# Genome Detective Coronavirus Typing Tool for rapid identification and characterization of novel coronavirus genomes

**DOI:** 10.1101/2020.01.31.928796

**Authors:** Sara Cleemput, Wim Dumon, Vagner Fonseca, Wasim Abdool Karim, Marta Giovanetti, Luiz Carlos Alcantara, Koen Deforche, Tulio de Oliveira

## Abstract

**Summary:** Genome Detective is a web-based, user-friendly software application to quickly and accurately assemble all known virus genomes from next generation sequencing datasets. This application allows the identification of phylogenetic clusters and genotypes from assembled genomes in FASTA format. Since its release in 2019, we have produced a number of typing tools for emergent viruses that have caused large outbreaks, such as Zika and Yellow Fever Virus in Brazil. Here, we present The Genome Detective Coronavirus Typing Tool that can accurately identify novel coronavirus (2019-nCoV) sequences isolated in China and around the world. The tool can accept up to 2,000 sequences per submission and the analysis of a new whole genome sequence will take approximately one minute. The tool has been tested and validated with hundreds of whole genomes from ten coronavirus species, and correctly classified all of the SARS-related coronavirus (SARSr-CoV) and all of the available public data for 2019-nCoV. The tool also allows tracking of new viral mutations as the outbreak expands globally, which may help to accelerate the development of novel diagnostics, drugs and vaccines.

**Availability:** Available online: https://www.genomedetective.com/app/typingtool/cov

**Supplementary information:** Supplementary data is available online.

## INTRODUCTION

We are currently faced with a potential global epidemic of a new coronavirus that has infected thousands of people in China and is spreading rapidly around the world. This week the WHO has declared it an global emergency (WHO, 2020). The Wuhan novel coronavirus (2019-nCoV) has already caused more infections than the previous severe acute respiratory syndrome (SARS) outbreak of 2002 and 2003. The virus is a SARS related coronavirus (SARSr-CoV), and it is genetically associated with SARSr-CoV strains that infect bats in China (Zhu et al., 2020, Lu et al., 2020). It causes severe respiratory illness, has high fatality rate (Huang et al., 2020), can be transmitted from person to person and has spread to over 15 countries in less than two months (WHO, 2020).

This coronavirus outbreak has been unprecedented; so too is the way that the scientific community has responded to it. They have openly and rapidly shared genomic and clinical data as never seen before allowing research results to be released almost instantaneously. This has helped the understanding of the transmission dynamics, the development of rapid diagnostic and has informed public health response. Here, we present a new contribution that can speed up this communal effort. The Genome Detective Coronavirus Typing Tool is a free-of-charge web-based bioinformatics pipeline that can accurately and quickly identify, assemble and classify coronaviruses genomes. The tool also identifies changes at nucleotides, coding regions and proteins using a novel dynamic aligner to allow tracking new viral mutations (Figure 1).

**Figure 1:**
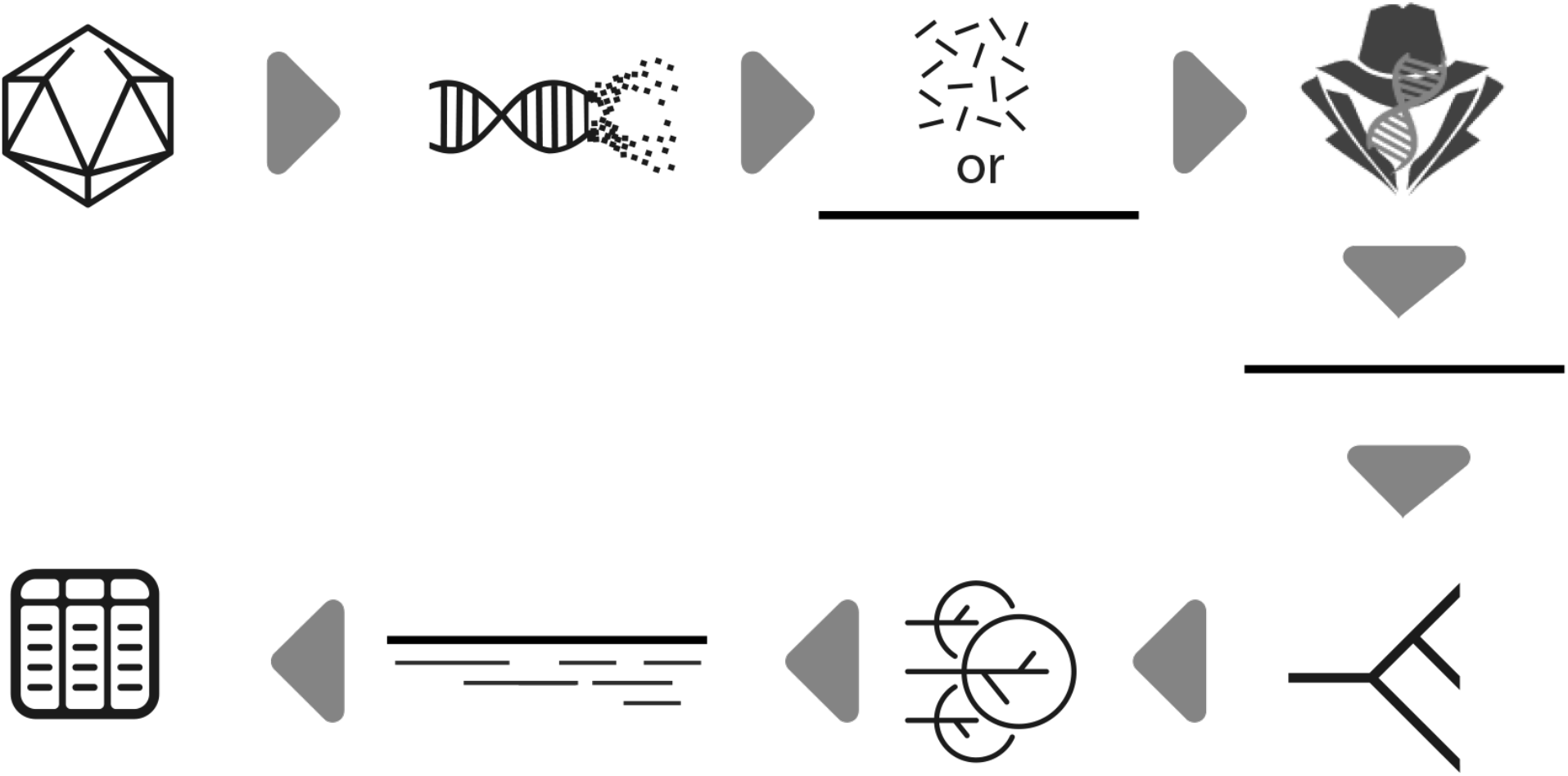
Genome Detective Coronavirus Typing Tool assembles genomes from next generation sequencing (NGS) in FASTAQ format or assembled genomes in FASTA format. A user can submit up to 1Gb of NGS data or 2,000 assembled genomic sequences. For each assembled genomic sequence, the tool identifies the virus species, constructs a phylogenetic tree and identifies phylogenetic clusters, which includes the novel coronavirus identified in Wuhan China in 2019 (2019-nCoV). The tool identifies changes at nucleotides, coding regions and proteins using a novel dynamic aligner and display all of the mutations in detailed tables and reports.

## SYSTEMS AND METHODS

A reference dataset of previously published coronavirus whole genome sequences (WGS) was compiled from the Virus Pathogen Resource (VIPR) database (www.viprbrc.org). This dataset consisted of 386 whole genome sequences (WGS) of nine important coronavirus species. These included 132 sequences of Severe Acute Respiratory Syndrome related Coronavirus (SARSr-CoV), 121 sequences of Beta coronavirus, 97 sequences of Middle East Respiratory Syndrome related Coronavirus (MERSr-CoV), 19 sequences of Human Coronavirus HKU1, 9 sequences of Murine Hepatitis Virus, 4 of Rousettus Bat Coronavirus HKU9, 3 of Rat Coronavirus and one WGS of Tylonycteris Bat Coronavirus HKU4, Zaria_bat_coronavirus and Longquan Rl Rat coronavirus). To this reference dataset, we added 47 whole genomes of the current Coronavirus 2019 (2019-nCoV) outbreak that originated in Wuhan, China, in December 2019. The 2019-nCoV sequences were downloaded from the GISAID database (https://www.gisaid.org) together with annotation of its original location, collection date and originating and submitting laboratory. The 2019-nCoV data generators are properly acknowledged in the acknowledgements section of this paper and detailed information is provided in Supplementary Table 1.

The 431 reference WGS were aligned with MUSCLE (Edgar 2004). The alignment was manually edited until a codon alignment was attained in all coding sequences (CDS). A Maximum likelihood phylogenetic tree, 1000 bootstrap replicates was constructed in PhyML (Guidon & Gascuel 2003; Lemoine et al., 2018) and a Bayesian tree using MrBayes (Ronquist & Huelsenbeck 2003) were constructed. The trees were visualized in Figtree (Rambaut 2018). We selected 25 reference sequences that represent the diversity of each well-defined phylogenetic cluster (with bootstrap support of 100% and posterior probability of 1). We identified five well supported phylogenetic clusters with more than two sequences of SARSr-CoV and used them to set up our automated phylogenetic classification tool. Cluster 1 included SARS strains from the 2002 and 2003 Asian outbreaks. In our tool, we named this cluster *SARS-CoV Outbreak 2000s but may rename it as SARS-A if a new proposed naming system for SARSr-Cov is adopted in the near future (Rambaut 2020)*. Cluster 2 (provisionally named as *SARS related CoV*) includes 7 sequences from bats which did not cause large human outbreaks. Cluster 3 (named as *Bat SARS-CoV HKU3*) includes three WGS sampled from Rhinolophus sinicus (i.e. Chinese rufous horseshoe bats). Cluster 4 (*Bat SARS-CoV ZXC21/ZC45*) includes two SARSr-CoV sampled from Rhinolophus sinicus bats in Zhoushan, China. Cluster 5 (provisionally named as *Wuhan 2019-nCoV, which may be renamed as SARS-B*) includes one public sequence from the outbreak in Wuhan, China. We identified this cluster with many sequences from GISAID but kept only this one as this is the only GenBank sequence, accession number MN908947, which was kindly shared by Prof. Yong-Zhen Zhang and colleagues in the virological.org website. Detailed information about the phylogenetic reference datasets are available in Supplementary Table 2.

The phylogenetic reference dataset was used to create an automated Coronavirus Typing Tool using the Genome Detective framework (Vilsker et al., 2019, Fonseca et al., 2019). To determine the accuracy of this tool, each of the 431 test WGS was considered for evaluation (i.e. 384 reference sequences from VIPR and 47 public 2019-nCoV sequences). The sensitivity, specificity and accuracy of our method was calculated for both species assignment and phylogenetic clustering of SARSr-CoV. Sensitivity was computed by the formula 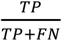, specificity by 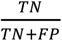 and accuracy by 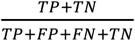, where TP = True Positives, FP = False Positives, TN = True Negatives and FN = False Negatives.

Classifying query sequences in an automated fashion involves two steps. The first step enables virus species assignments and the second, which is restricted to SARSr-CoV, includes phylogenetic analysis. The first classification analysis subjects a query sequence to BLAST and AGA analysi. AGA is a novel alignment method for nucleic acid sequences against annotated genomes from NCBI RefSeq Virus Database. AGA (Deforche 2017) expands the optimal alignment algorithms of Smith-Waterman (Smith & Waterman 1981) and Gotoh (Gotoh 1982) based on an induction state with additional parameters. The result is a more accurate aligner, as it takes into account both nucleotide and protein scores and identifies all of the polymorphisms at nucleotide and amino acid levels. In the second step, a query sequence is aligned against the phylogenetic reference dataset using -add alignment option in the MAFFT software (Katoh & Standley 2013). In addition, a Neighbor Joining phylogenetic tree is constructed using the HKY distance metric with gamma among-site rate variation with 1,000 bootstrap replicates using PAUP* (Swofford). The query sequence is assigned to a particular phylogenetic cluster if it clusters monophyletically with that clade or a subset of it with bootstrap support >70%. If the bootstrap support is <70%, the genotype is reported to be unassigned.

The result of the phylogenetic and mutational analysis performed by AGA is available in a detailed report. This report contains an interactive phylogenetic tree and genome mapper (Supplementary Figure 1). It also presents the virus species and cluster assignments and a detailed table that provides information about open reading frames (ORFs), CDS and proteins. This table can be expanded to show nucleotide and amino acid mutations that differentiate a query sequence from their species RefSeq or from a sequence in the phylogenetic reference dataset. All results can be exported to a variety of file formats (XML, CSV, Excel, Nexus or FASTA).

## TESTING AND VALIDATION

The Genome Detective Coronavirus Typing Tool correctly classified all of the 175 SARSr-CoV sequences at species level, i.e. specificity, sensitivity and accuracy of 100%. Furthermore, all of the 47 2019-nCoV WGS that were isolated in China (n=36), USA (n=5), France (n=2), Thailand (n=2), Japan (n=1) and Taiwan (n=1) were correctly classified at phylogenetic cluster level as 2019-nCoV, which may be renamed as SARS-B. In addition, we classified with very high specificity, sensitivity and accuracy (i.e. 100%) all of the 112 SARS outbreak WGS of 2002 and 2003. We also achieved perfect classification (i.e. specificity, sensitivity and accuracy of 100%) for all of Beta coronavirus, Human_coronavirus_HKU1, MERS-CoV, Rousettus_bat_coronavirus_HKU9 and Tylonycteris_bat_coronavirus_HKU4 at species level. For a detailed overview of assignment performance, please refer to the Supplementary Table 3.

Our tool also allows detailed analysis of coding regions and proteins for each of the coronavirus species. For example, the analysis of the first released 2019-nCoV sequence, the WH_Human1_China_2019Dec (GenBank: MN908947) demonstrated at genome level, the nucleotide (NT) identity was 79.0% to the reference strain of SARSr-CoV (ACCESSION: NC_004718.3) and that the Envelop Small Membrane Protein (protein E) is the most similar protein. In total, 94.8% (73/77) of the amino acids were identical; the four amino acid differences were located at positions 55 (T55S), 56 (V56F), 69 (69deletion) and 70 (G70R). The spike protein (protein S), which can be associated with virulence, was 76.2% identical to the reference strain of SARSr-CoV (Supplementary Table 4A). Interestingly there were four amino acid insertions at position 237 (A237_F238insHRSY, genome NT position 22202_22203insCATAGAAGTTAT)), which is just upstream from a cleavage site. The most diverse coding regions were the CDS Sars8a and Sars8b. In these two regions, only 30% of the amino acids were identical. Sars8b protein was truncated early and its CDS had four stop codons (Supplementary Table 4sA).

Our Coronavirus Typing Tool also allows a query sequence to be analysed against a sequence in the phylogenetic reference dataset. For example, the WH_Human1_China_2019Dec (GenBank: MN908947) the identity was 87.5% to the Bat sequence bat_SL_CoVZXC21 (Genbank: MG772934). This was one of the Bat-CoV sequences that were most related to n2019-CoV (Lu et al. 2020). The Envelop Small Membrane Protein (protein E) was 100% identical (Supplementary Table 4B). When the 2019-nCoV isolated from France (BetaCoV/France/IDF0373/2020) was analysed with our tool and compared with the 2019-nCOv WH_Human1_China_2019Dec strain (Accession: MN908947), this sequence was 99.9% identical and had only two NT mutations (Supplementary Table 4C). These two differences were located on positions: 22551G>T & 26016G>T), which caused three amino acid mutations (E2 glycoprotein Protein mutation: V354F (22551G>T), sars3a protein mutations: G250V (26016G>T) and sars3b protein mutations: V110F (26016G>T)) (Detailed in Supplementary Table 4C-II). The analysis of a WGS in FASTA format takes approximately 60 seconds.

## DISCUSSION

We developed and released the Genome Detective Coronavirus Typing tool as a free-of-charge resource in the third week of January 2020 in order to help the rapid characterization of nCoV-2019 infections. This tool allows the analysis of whole or partial viral genomes within minutes. It accepts assembled genomes in FASTA format or raw NGS data in FASTQ format from Illumina, Ion Torrent, PACBIO or Oxford Nanopore Technologies (ONT) can be submitted to the Genome Detective Virus Tool (Vilsker et al., 2019) to automatically assemble the consensus genome prior to executing the Coronavirus Typing Tool. User effort is minimal, and a user can submit multiple FASTA sequences at once.

The tool uses a novel and dynamic aligner, AGA, to allow submitted sequences to be queried against reference genomes, using both nucleotide and amino acid similarity scores. This allows accurate identification of other coronavirus species and the tracking of new viral mutations as the outbreak expands globally. It also performs detailed analysis of the coding regions and proteins. Moreover, it can easily be updated to add new phylogenetic clusters if new outbreaks arise or if the classification nomenclature changes. The tool has been able to correctly classify all the recently released nCoV-2019 genomes, as well as all the 2002-2003 SARS outbreak sequences

In conclusion, the Genome Detective Coronavirus Typing Tool is a web-based and user-friendly software application that allows the identification and characterization of novel coronavirus genomes.

## ACKNOWLEDGEMENTS

Genome data for 2019-nCoV made kindly available by National Institute for Communicable Disease Control and Prevention (ICDC) Chinese Center for Disease Control and Prevention (China CDC), Institute of Pathogen Biology, Chinese Academy of Medical Sciences & Peking Union Medical College, Hubei Provincial Center for Disease Control and Prevention, Wuhan Institute of Virology, Chinese Academy of Sciences, National Institute for Viral Disease Control and Prevention, China CDC, Li Ka Shing Faculty of Medicine, The University of Hong Kong, Department of Medical Sciences, Ministry of Public Health, Thailand | Thai Red Cross Emerging Infectious Diseases - Health Science Centre | Department of Disease Control, Ministry of Public Health, Thailand, Department of Microbiology, Guangdong Provincial Center for Diseases Control and Prevention, Department of Microbiology, Zhejiang Provincial Center for Disease Control and Prevention, Division of Viral Diseases, Centers for Disease Control and Prevention, Centers for Disease Control, R.O.C. (Taiwan) | Centers for Disease Control, R.O.C. (Taiwan), California Department of Public Health | Pathogen Discovery, Respiratory Viruses Branch, Division of Viral Diseases, Centers for Diseases Control and Prevention, Arizona Department of Health Services | Pathogen Discovery, Respiratory Viruses Branch, Division of Viral Diseases, Centers for Disease Control and Prevention, Guangdong Provincial Centers for Diseases Control and Prevention | Guangdong Provincial Institute of Public Health, University of Hong Kong-Shenzhen Hospital, Shenzhen, Guangdong, State Key Laboratory of Virology, Wuhan University, Department of Infectious and Tropical Diseases, Bichat Claude Bernard Hospital, Paris | National Reference Center for Viruses of Respiratory Infections, Institut Pasteur, Paris. We would like to acknowledge all of the data contributors. Research reported in this publication was supported by a research Flagship grant from the South African Medical Research Council (MRC-RFA-UFSP-01-2013/UKZN HIVEPI), the VIROGENESIS project, which received funding from the European Union’s Horizon 2020 Research and Innovation Programme (under Grant Agreement no. 634650) and the National Human Genome Research Institute of the National Institutes of Health under Award Number U24HG006941. H3ABioNet is an initiative of the Human Health and Heredity in Africa Consortium (H3Africa). The content is solely the responsibility of the authors and does not necessarily represent the official views of the National Institutes of Health.

## REFERENCES

Zhu N, Zhang D, Wang W, Li X, Yang B, Song J, Zhao X, Huang B, Shi W, Lu R, Niu P, Zhan F, Ma X, Wang D, Xu W, Wu G, Gao GF, Tan W; China Novel Coronavirus Investigating and Research Team. A Novel Coronavirus from Patients with Pneumonia in China, 2019. N Engl J Med. 2020, doi: 10.1056/NEJMoa2001017.

Huang C, Wang Y, Li X, Ren L, Zhao J, Hu Y, Zhang L, Fan G, Xu J, Gu X, Cheng Z, Yu T, Xia J, Wei Y, Wu W, Xie X, Yin W, Li H, Liu M, Xiao Y, Gao H, Guo L, Xie J, Wang G, Jiang R, Gao Z, Jin Q, Wang J, Cao B. Clinical features of patients infected with 2019 novel coronavirus in Wuhan, China. Lancet. 2020, pii: S0140-6736(20)30183–5. doi: 10.1016/S0140-6736(20)30183-5.

Lu R, Xiang Zhao, Juan Li, Peihua Niu, Bo Yang, Honglong Wu, Wenling Wang, Hao Song, Baoying Huang, Na Zhu, Yuhai Bi, Xuejun Ma, Faxian Zhan, Liang Wang, Tao Hu, Hong Zhou, Zhenhong Hu, Weimin Zhou, Li Zhao, Jing Chen, Yao Meng, Ji Wang, Yang Lin, Jianying Yuan, Zhihao Xie, Jinmin Ma, William J Liu, Dayan Wang, Wenbo Xu, Edward C Holmes, George F Gao, Guizhen Wu, Weijun Chen, Weifeng Shi, Wenjie Tan. Genomic characterisation and epidemiology of 2019 novel coronavirus: implications for virus origins and receptor binding. Lancet. 2020. DOI:https://doi.org/10.1016/S0140-6736(20)30251-8

World Health Organization (WHO) Novel Coronavirus (2019-nCoV) situational reports. Available online https://www.who.int/emergencies/diseases/novel-coronavirus-2019/situation-reports/

Edgar RC. MUSCLE: multiple sequence alignment with high accuracy and high throughput. Nucleic Acids Research. 2004, 32(5): 1792–97. doi:10.1093/nar/gkh340

Guindon S, Gascuel O. A simple, fast, and accurate algorithm to estimate large phylogenies by maxi- mum likelihood. Systematic Biology. 2003; 52(5):696–704. https://doi.org/10.1080/10635150390235520

Lemoine F, Domelevo Entfellner JB, Wilkinson E, Correia D, Davila Felipe M, de Oliveira T, Gascuel O. Renewing Felsenstein’s phylogenetic bootstrap in the era of big data. Nature. 2018, doi:10.1038/s41586-018-0043-0

Ronquist F, Huelsenbeck JP. MrBayes 3: Bayesian phylogenetic inference under mixed models. Bioin- formatics. 2003; 19(12):1572–1574. https://doi.org/10.1093/bioinformatics/btg180

Rambaut A. FigTree v1.4.4. Institute of Evolutionary Biology, University of Edinburgh, Edinburgh. 2018, http://tree.bio.ed.ac.uk/software/figtree/

Rambaut A. Twitter, 24 Jan 2020. https://twitter.com/arambaut

Vilsker M, Moosa Y, Nooij S, Fonseca V, Ghysens Y, Dumon K, Pauwels R, Alcantara LC, Vanden Eynden E, Vandamme AM, Deforche K, de Oliveira T. Genome Detective: an automated system for virus identification from high-throughput sequencing data. Bioinformatics. 2019; 35(5):871–873. doi: 10.1093/bioinformatics/bty695.

Fonseca V, Libin PJK, Theys K, Faria NR, Nunes MRT, Restovic MI, Freire M, Giovanetti M, Cuypers L, Nowé A, Abecasis A, Deforche K, Santiago GA, Siqueira IC, San EJ, Machado KCB, Azevedo V, Filippis AMB, Cunha RVD, Pybus OG, Vandamme AM, Alcantara LCJ, de Oliveira T. A computational method for the identification of Dengue, Zika and Chikungunya virus species and genotypes. PLoS Negl Trop Dis. 2019; 13(5):e0007231. doi: 10.1371/journal.pntd.0007231.

Deforche K. An alignment method for nucleic acid sequences against annotated genomes. 2017. doi.org/10.1101/200394.

Smith T.F., Waterman M.S. (1981) Identification of common molecular subsequences. J. Mol. Biol., 147, 195–197.

Gotoh O. (1982) An improved algorithm for matching biological sequences. J. Mol. Biol., 162, 705–708.

Katoh K, Standley DM. MAFFT multiple sequence alignment software version 7: Improvements in performance and usability. Molecular Biology and Evolution. 2013;30(4):772–780.

Swofford,D.L. PAUP* 4.0: phylogenetic analysis under parsimony (and other methods), version 4.0b2a. Sinauer Associates Inc., Sunderland, Mass.

