## Supplementary Figure 1 for "Genome Detective Coronavirus Typing Tool for rapid identification and characterization of novel coronavirus genomes"

NO JOBS IN QUEUE

### Coronavirus Typing Tool

#### CORONAVIRUS TYPING TOOL DETAILS

Version 1.9

##### SEQUENCE ASSIGNMENT

**Name** MN908947.3

**Length** 29820

###### VIRUS ASSIGNMENT

**Virus assignment** Severe acute respiratory syndrome-related coronavirus

###### CLADE AND GENOTYPE RESULT

**Clade assignment** Wuhan 2019-nCoV

Supported with phylogenetic analysis and bootstrap 100.0 (>= 70.0)

###### GENOME REGION

Sequence starts at position 1 and ends at position 29677 relative to the NC_004718.3 reference sequence for Severe acute respiratory syndrome-related coronavirus (taxon:694009).

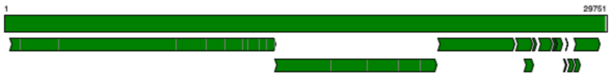


##### PHYLOGENETIC ANALYSIS DETAILS (GENOTYPE)

- Assignment: Wuhan 2019-nCoV
- Bootstrap support: 100.0
- Download the alignment (NEXUS format, FASTA format)
- Phylogenetic Tree (export as PDF, NEXUS Format)

**TREE CONTROLS**

**Layout**

Rectilinear

**Transform**

None

- Show labels
- Highlight clusters
- Color branches

Bat SARS-CoV ZXC21/ZC45

Wuhan 2019-nCoV

Bat SARS-CoV HKU3

SARS related CoV

SARS-CoV Outbreak 2000s

outgroup

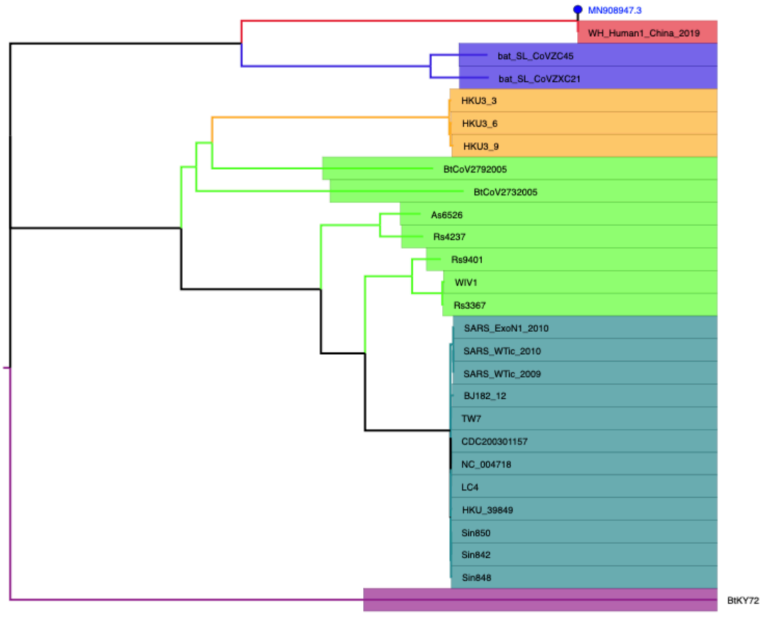

#### Phylogenetic tree can be visualized in polar or radial format:

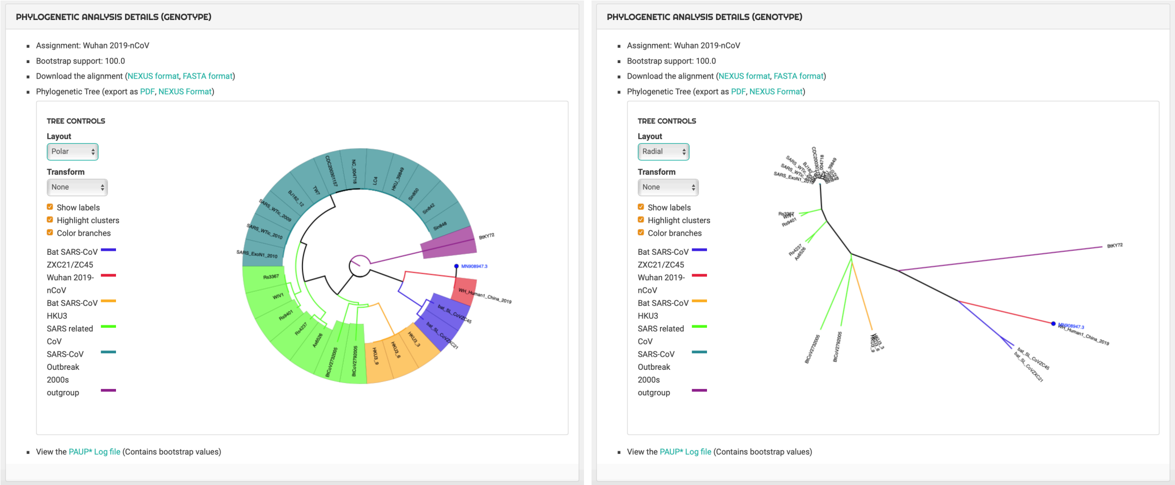


---

#### Alignment Details and Genome Mapper:

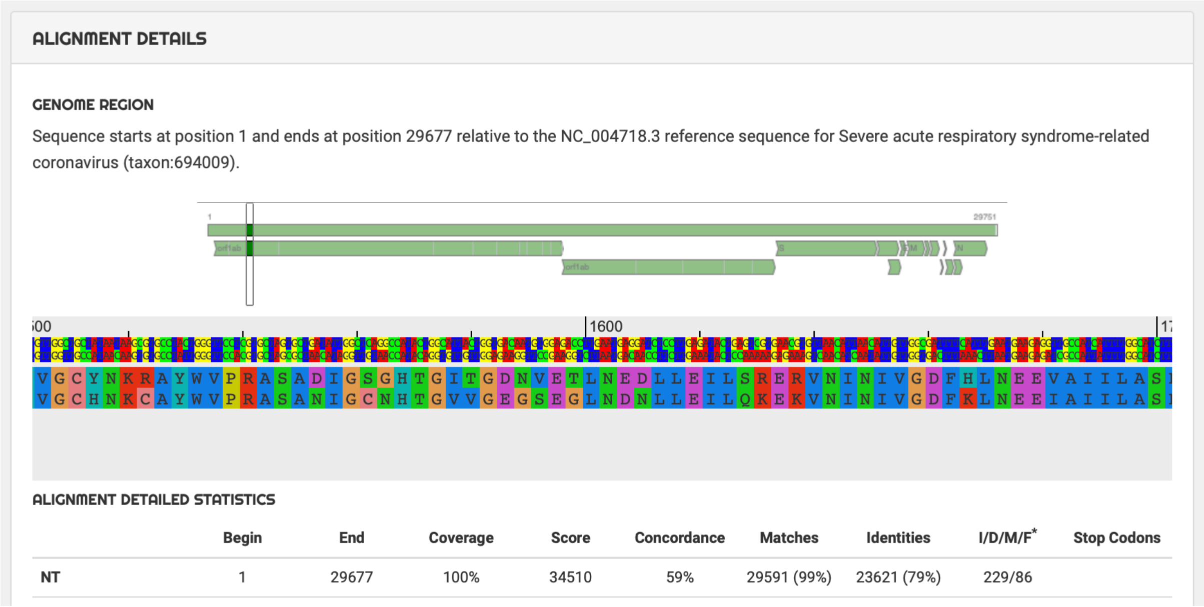

ALIGNMENT DETAILS

GENOME REGION

Sequence starts at position 1 and ends at position 29677 relative to the NC_004718.3 reference sequence for Severe acute respiratory syndrome-related coronavirus (taxon:694009).



ALIGNMENT DETAILED STATISTICS

| | Begin | End | Coverage | Score | Concordance | Matches | Identities | I/D/M/F* | Stop Codons |
|---|---|---|---|---|---|---|---|---|---|
| NT | 1 | 29677 | 100% | 34510 | 59% | 29591 (99%) | 23621 (79%) | 229/86 | |

#### Nucleotide (NT) and Coding Regions (CDS) Mutational Analysis

SHOW MUTATIONS

| | Begin | End | Coverage | Score | Concordance | Matches | Identities | I/D/M/F* | Stop Codons |
|---|---|---|---|---|---|---|---|---|---|
| **NT** | 1 | 29677 | 99.8% | 34510 | 58.6% | 29591 (98.9%) | 23621 (79.0%) | 229/86 | |
| **CDS** | | | | | | | | | |
| 1_orf1ab | 1 | 7074 | 100% | 44034 | 89.0% | 7068 (99.5%) | 6126 (86.2%) | 29/6/0/0 | 1 |
| 2_orf1ab | 1 | 4383 | 100% | 25297 | 83.9% | 4377 (99.2%) | 3553 (80.5%) | 29/6/0/0 | 1 |
| 3_S | 1 | 1256 | 100% | 7176 | 81.6% | 1249 (97.5%) | 976 (76.2%) | 25/7/0/0 | 1 |
| 4_sars3a | 1 | 275 | 100% | 1490 | 75.5% | 275 (99.6%) | 200 (72.5%) | 1/0/0/0 | 1 |
| 5_sars3b | 1 | 155 | 100% | 519 | 52.9% | 155 (99.4%) | 89 (57.1%) | 1/0/0/0 | 5 |
| 6_E | 1 | 77 | 100% | 447 | 96.3% | 76 (98.7%) | 73 (94.8%) | 0/1/0/0 | 1 |
| 7_M | 1 | 222 | 100% | 1419 | 92.9% | 222 (99.6%) | 202 (90.6%) | 1/0/0/0 | 1 |
| 8_sars6 | 1 | 64 | 100% | 300 | 74.6% | 62 (96.9%) | 43 (67.2%) | 0/2/0/0 | 1 |
| 9_sars7a | 1 | 123 | 100% | 758 | 89.3% | 122 (99.2%) | 105 (85.4%) | 0/1/0/0 | 1 |
| 10_sars7b | 1 | 45 | 100% | 252 | 83.7% | 44 (97.8%) | 36 (80.0%) | 0/1/0/0 | 1 |
| 11_sars8a | 1 | 40 | 100% | 100 | 33.8% | 39 (90.7%) | 13 (30.2%) | 3/1/0/0 | 0 |
| 12_sars8b | 1 | 85 | 100% | 26 | 4.2% | 81 (81.8%) | 30 (30.3%) | 14/4/1/1 | 4 |
| 13_N | 1 | 423 | 100% | 2645 | 91.3% | 420 (99.3%) | 383 (90.5%) | 0/3/0/0 | 1 |
| 14_sars9b | 1 | 99 | 100% | 451 | 72.9% | 98 (99.0%) | 72 (72.7%) | 0/1/0/0 | 1 |

#### Protein Analysis (from UNIPROT RefSeq):

| **Proteins** | | | | | | | | | |
|---|---|---|---|---|---|---|---|---|---|
| orf1ab polyprotein... | 1 | 7074 | 100% | 44034 | 89.0% | 7068 (99.5%) | 6126 (86.2%) | 29/6/0/0 | 1 |
| leader protein (NP... | 1 | 180 | 100% | 1059 | 87.4% | 180 (100%) | 152 (84.4%) | 0/0/0/0 | 0 |
| counterpart of MH... | 1 | 638 | 100% | 3197 | 71.2% | 638 (100%) | 436 (68.3%) | 0/0/0/0 | 0 |
| nsp3-pp1a/pp1ab ... | 1 | 1922 | 100% | 10415 | 80.4% | 1916 (98.2%) | 1486 (76.2%) | 29/6/0/0 | 0 |
| nsp4-pp1a/pp1ab ... | 1 | 500 | 100% | 3038 | 85.4% | 500 (100%) | 400 (80.0%) | 0/0/0/0 | 0 |
| 3C-like proteinase ... | 1 | 306 | 100% | 2172 | 97.4% | 306 (100%) | 294 (96.1%) | 0/0/0/0 | 0 |
| nsp6-pp1a/pp1ab ... | 1 | 290 | 100% | 1813 | 89.9% | 290 (100%) | 253 (87.2%) | 0/0/0/0 | 0 |
| nsp7-pp1a/pp1ab ... | 1 | 83 | 100% | 508 | 98.6% | 83 (100%) | 82 (98.8%) | 0/0/0/0 | 0 |
| nsp8-pp1a/pp1ab ... | 1 | 198 | 100% | 1210 | 97.6% | 198 (100%) | 193 (97.5%) | 0/0/0/0 | 0 |
| nsp9-pp1a/pp1ab ... | 1 | 113 | 100% | 752 | 96.9% | 113 (100%) | 110 (97.3%) | 0/0/0/0 | 0 |
| formerly known as... | 1 | 139 | 100% | 1061 | 97.7% | 139 (100%) | 135 (97.1%) | 0/0/0/0 | 0 |
| RNA-dependent R... | 1 | 932 | 100% | 6561 | 97.1% | 932 (100%) | 898 (96.4%) | 0/0/0/0 | 0 |
| nsp13-pp1ab (ZD, ... | 1 | 601 | 100% | 4241 | 99.9% | 601 (100%) | 600 (99.8%) | 0/0/0/0 | 0 |
| 3'-to-5' exonucleas... | 1 | 527 | 100% | 3864 | 96.5% | 527 (100%) | 501 (95.1%) | 0/0/0/0 | 0 |
| endoRNAse (NP_8... | 1 | 346 | 100% | 2174 | 92.6% | 346 (100%) | 307 (88.7%) | 0/0/0/0 | 0 |
| 2'-O-ribose methylt... | 1 | 298 | 100% | 1968 | 95.4% | 298 (100%) | 278 (93.3%) | 0/0/0/0 | 0 |
| orf1a polyprotein (... | 1 | 4383 | 100% | 25297 | 83.9% | 4377 (99.2%) | 3553 (80.5%) | 29/6/0/0 | 1 |
| nsp11-pp1a (NP_9... | 1 | 13 | 100% | 71 | 89.9% | 13 (100%) | 11 (84.6%) | 0/0/0/0 | 0 |
| E2 glycoprotein pr... | 1 | 1256 | 100% | 7176 | 81.6% | 1249 (97.5%) | 976 (76.2%) | 25/7/0/0 | 1 |

#### 2019-nCov Protein Mutations and Codon Mutations on E and Matrix Proteins as compared with SARSr-CoV RefSeq (GenBank: NC_004718.3 ).

| | | | | | | | | | |
|---|---|---|---|---|---|---|---|---|---|
| protein E (NP_828… | 1 | 77 | 100% | 447 | 96.3% | 76 (98.7%) | 73 (94.8%) | 0/1/0/0 | 1 |
| Protein mutations: | T55S (26279A>T 26281G>T), V56F (26282G>T), E69del (26321_26323delGAA), G70R (26324G>A) | | | | | | | | |
| Codon mutations: | GAA8GAG (26140A>G), GTC29GTT (26203C>T), TTA51CTT (26267T>C 26269A>T), CCA54CCT (26278A>T), ACG55TCT (26279A>T 26281G>T), GTT56TTT (26282G>T), GTC58GTT (26290C>T), TCG60TCT (26296G>T), AAC66AAT (26314C>T), GAA69del (26321_26323delGAA), GGA70AGA (26324G>A) | | | | | | | | |
| matrix protein (NP… | 1 | 222 | 100% | 1419 | 92.9% | 222 (99.6%) | 202 (90.6%) | 1/0/0/0 | 1 |
| Protein mutations: | D3_N4insS (26405_26406insTTC), Q14K (26437C>A 26439A>G), A29T (26482G>A 26484C>A), M32C (26491A>T 26492T>G 26493G>T), S39A (26512T>G 26514T>C), V51I (26548G>A), V75I (26620G>A 26622G>C), I86L (26653A>C), V96I (26683G>A), R124H (26768G>A 26769G>T), V128L (26779G>C), M133L (26794A>C 26796G>A), I144L (26827A>C), M150I (26847G>T), S154H (26857T>C 26858C>A 26859C>T), G187A (26957G>C 26958C>A), T188G (26959A>G 26960C>G), N196S (26984A>G 26985C>T), A210S (27025G>T), G211S (27028G>A), N213S (27035A>G 27036C>T) | | | | | | | | |
| Codon mutations: | GAC3GAT (26405_26406insTTC), GAC3_AAC4insTCC (26405_26406insTTC), GAG10GAA (26427G>A), CAA14AAG (26437C>A 26439A>G), CTG16CTT (26445G>T), CTA28CTT (26481A>T), GCC29ACA (26482G>A 26484C>A), ATG32TGT (26491A>T 26492T>G 26493G>T), TTA33CTT (26494T>C 26496A>T), TCT39GCC (26512T>G 26514T>C), AAT40AAC (26517T>C), CGG41AGG (26518C>A), AAC42AAT (26523C>T), TAC46TAT (26535C>T), ATA48ATT (26541A>T), CTT50TTA (26545C>T 26547T>A), GTT51ATT (26548G>A), CTC55CTG (26562C>G), TTG56TTA (26565G>A), ACA60ACT (26577A>T), CTT61TTA (26578C>T 26580T>A), GTC69GTT (26604C>T), ATT72ATA (26613T>A), GTG75ATC (26620G>A 26622G>C), ACT76ACC (26625T>C), GGC77GGT (26628C>T), GGG78GGA (26631G>A), GCG80GCT (26637G>T), ATT81ATC (26640T>C), ATT86CTT (26653A>C), CTT92CTC (26673T>C), GTT96ATT (26683G>A), TCC98TCT (26691C>T), AGG100AGA (26697G>A), GCT103GCG (26706T>G), ACC105ACG (26712C>G), CGC106CGT (26715C>T), TCA107TCC (26718A>C), AAC112AAT (26733C>T), ACA115ACT (26742A>T), AAT120AAC (26757T>C), CCT122CCA (26763T>A), CGG124CAT (26768G>A 26769G>T), GGG125GGC (26772G>C), ACA126ACT (26775A>T), GTG128CTG (26779G>C), CTC132CTT (26793C>T), ATG133CTA (26794A>C 26796G>A), CTT137CTC (26808T>C), GTC138GTA (26811C>A), ATT139ATC (26814T>C), GGT140GGA (26817T>A), ATT144CTT (26827A>C), GGT146GGA (26835T>A), CAC147CAT (26838C>T), TTG148CTT (26839T>C 26841G>T), CGA149CGT (26844A>T), ATG150ATT (26847G>T), GCC151GCT (26850C>T), TCC154CAT (26857T>C 26858C>A 26859C>T), GGG156GGA (26865G>A), ATT160ATC (26877T>C), CCA164CCT (26889A>T), GAG166GAA (26895G>A), GTG169GTT (26904G>T), TTA180TTG (26937A>G), GCG182GCT (26943G>T), GGC187GCA (26957G>C 26958C>A), ACT188GGT (26959A>G 26960C>G), GAT189GAC (26964T>C), AAC196AGT (26984A>G 26985C>T), CGT199AGG (26992C>A 26994T>G), GGA201GGC (27000A>C), AAT206AAC (27015T>C), CAC209CAT (27024C>T), GCC210TCC (27025G>T), GGT211AGT (27028G>A), AAC213AGT (27035A>G 27036C>T), CTA219CTT (27054A>T) | | | | | | | | |

#### 2019-nCov Protein Mutations and Codon Mutations on E and Matrix Proteins as compared with Bat SARS related CoV sequence, bat_SL_CovZXC21 (GenBank: MG772934)

| | | | | | | | | | |
|---|---|---|---|---|---|---|---|---|---|
| protein E (NP_828… | 1 | 77 | 98.7% | 474 | 100% | 76 (100%) | 76 (100%) | 0/0/0/0 | 1 |
| Protein mutations: | *none* | | | | | | | | |
| Codon mutations: | TTT23TTC (26185T>C), GTC29GTT (26203C>T), TTG75CTG (26339T>C) | | | | | | | | |
| matrix protein (NP… | 1 | 223 | 100% | 1521 | 98.6% | 223 (100%) | 220 (98.7%) | 0/0/0/0 | 1 |
| Protein mutations: | S2A (26401T>G), G3D (26405G>A), D3i1S (26405i2G>T 26405i3A>C) | | | | | | | | |
| Codon mutations: | TCA2GCA (26401T>G), GGT3GAT (26405G>A), GAC3i1TCC (26405i2G>T 26405i3A>C), ACC6ACT (26415C>T), TTA16CTT (26443T>C 26445A>T), GGA24GGT (26469A>T), TTG26CTA (26473T>C 26475G>A), TTT27TTC (26478T>C), TTG33CTT (26494T>C 26496G>T), TTA34CTA (26497T>C), TAC46TAT (26535C>T), CTT56TTA (26563C>T 26565T>A), TGC63TGT (26586C>T), AAC73AAT (26616C>T), ACT76ACC (26625T>C), GCC80GCT (26637C>T), ATT81ATC (26640T>C), GCC84GCT (26649C>T), CTT92CTC (26673T>C), AGG100AGA (26697G>A), GCT103GCG (26706T>G), TTT111TTC (26730T>C), AAC112AAT (26733C>T), TTG119CTC (26752T>C 26754G>C), CCT122CCA (26763T>A), CTT123CTC (26766T>C), ACA126ACT (26775A>T), AGG130AGA (26787G>A), GAG134GAA (26799G>A), ATT139ATC (26814T>C), GCA151GCT (26850A>T), CTG155CTA (26862G>A), CCC164CCT (26889C>T), GTA169GTT (26904A>T), GGT201GGC (27000T>C), AAT202AAC (27003T>C), TAC203TAT (27006C>T), AAT206AAC (27015T>C) | | | | | | | | |

### 2019-nCov comparison between sequences isolated in France, Jan 2020 (BetaCoV/France/IDF0373/2020) and Wuhan, China< Dec 2019 (GenBank: MN908947)

#### ALIGNMENT DETAILS

##### ALIGNMENT

Using **NC_004718.3** (Severe acute respiratory syndrome-related coronavirus (taxon:694009)) as reference for alignment, numbering and genome annotations.

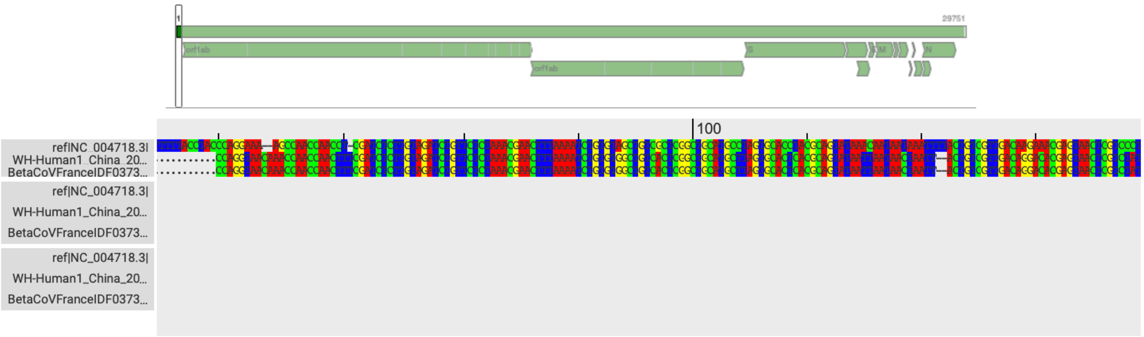


##### GENETIC DIVERSITY ANALYSIS

Showing genetic similarity and mutations of the sequence against a reference of choice.

**Reference for genetic diversity:** WH-Human1_China_2019-Dec

HIDE MUTATIONS

| | Begin | End | Coverage | Score | Concordance | Matches | Identities | I/D/M/F* | Stop Codons |
|---|---|---|---|---|---|---|---|---|---|
| **NT** | 20 | 29914 | 99.4% | 59610 | 99.9% | 29809 (100%) | 29807 (99.9%) | 0/0 | |
| Mutations: | 22551G>T, 26016G>T | | | | | | | | |

| | | | | | | | | | |
|---|---|---|---|---|---|---|---|---|---|
| E2 glycoprotein pr... | 1 | 1281 | 99.5% | 8985 | 99.9% | 1274 (100%) | 1273 (99.9%) | 0/0/0/0 | 1 |
| Protein mutations: | V354F (22551G>T) | | | | | | | | |
| Codon mutations: | GTC354TTC (22551G>T) | | | | | | | | |
| hypothetical protei... | 1 | 276 | 100% | 1944 | 99.4% | 276 (100%) | 275 (99.6%) | 0/0/0/0 | 1 |
| Protein mutations: | G250V (26016G>T) | | | | | | | | |
| Codon mutations: | GGT250GTT (26016G>T) | | | | | | | | |
| hypothetical protei... | 1 | 156 | 100% | 964 | 99.6% | 156 (100%) | 155 (99.4%) | 0/0/0/0 | 5 |
| Protein mutations: | V110F (26016G>T) | | | | | | | | |
| Codon mutations: | GTT110TTT (26016G>T) | | | | | | | | |
| protein E (NP_828... | 1 | 77 | 98.7% | 474 | 100% | 76 (100%) | 76 (100%) | 0/0/0/0 | 1 |

Submission page: Input sequences in FASTA format

##### CORONAVIRUS TYPING TOOL

This tool is designed to use Blast and phylogenetic methods in order to identify the Coronavirus types and genotypes of a nucleotide sequence.

The Coronavirus typing tool also includes a Wuhan Coronavirus genome, this sequence was generously shared by Professor Yong-Zhen Zhang and colleagues via this post on virological.org. Professor Yong-Zhen Zhang and colleagues ask that you communicate with them if you wish to publish results that use the sequence that they share in a journal. We gratefully acknowledge their contribution.

**Note for batch analysis:** The tool accepts up to 2000 sequences at a time.

##### INPUT

Submit one or more FASTA sequences to be typed individually. If you have raw NGS reads (short reads or long reads), please use the Genome Detective Virus Tool to assemble first. Subtyping tools will be linked in the results.
Click here to load some sample data.

**Sequence**

CLICK OR DROP FILE

```
>MN908947.3 Wuhan seafood market pneumonia
virus isolate Wuhan-Hu-1, complete genome
ATTAAAGGTTTATACCTTCCCAGGTAACAAACCAACCAACTTTCGAT
CTCTTGTAGATCTGTTCTCTAAACGAACTTTAAAATCTGTGTGGCTG
TCACTCGGCTGCATGCTTAGTGCACTCACGCAGTATAATTAATAACT
AATTACTGTCGTTGACAGGACACGAGTAACTCGTCTATCTTCTGCAG
GCTGCTTACGGTTTCGTCCGTGTTGCAGCCGATCATCAGCACATCTA
GGTTTCGTCCGGGTGTGACCGAAAGGTAAGATGGAGAGCCTTGTCCC
TGGTTTCAACGAGAAAACACACGTCCAACTCAGTTTGCCTGTTTTAC
AGGTTCGCGACGTGCTCGTACGTGGCTTTGGAGACTCCGTGGAGGAG
GTCTTATCAGAGGCACGTCAACATCTTAAAGATGGCACTTGTGGCTT
AGTAGAAGTTGAAAAAGGCGTTTTGCCTCAACTTGAACAGCCCTATG
TGTTCATCAAACGTTCGGATGCTCGAACTGCACCTCATGGTCATGTT
ATGGTTGAGCTGGTAGCAGAACTCGAAGGCATTCAGTACGGTCGTAG
TGGTGAGACACTTGGTGTCCTTGTCCCTCATGTGGGCGAAATACCAG
[illegible]
```

START FREE ANALYSIS    CLEAR

Log in or register to experience the advantages of a premium account.
