## Supplementary Tables 1 to 4 for "Genome Detective Coronavirus Typing Tool for rapid identification and characterization of novel coronavirus genomes"

**Supplementary Table 1:** n2019-CoV GISAID Tested Sequences. GISAID acknowledges the Authors and the Laboratories for their sequence and metadata shared through GISAID. All Originating laboratories are gratefully acknowledged for sharing the data and are cited in this table:

| GISAID Accession ID | Virus name | Location | Collection date | Lab citation |
|---|---|---|---|---|
| EPI_ISL_402125 | BetaCoV/Wuhan-Hu-1/2019 | China | 2019-12 | 1 |
| EPI_ISL_405839 | BetaCoV/Guangdong/20SF013/2031 | China / Guangdong / Shenzhen | 2020-01 | 2 |
| EPI_ISL_406030 | BetaCoV/Guangdong/20SF013/2032 | China / Guangdong / Shenzhen | 2020-01 | 2 |
| EPI_ISL_402119 | BetaCoV/Wuhan/IVDC-HB-01/2019 | China / Hubei Province / Wuhan City | 2019-12-30 | 3 |
| EPI_ISL_402120 | BetaCoV/Wuhan/IVDC-HB-04/2020 | China / Hubei Province / Wuhan City | 2020-01-01 | 3 |
| EPI_ISL_402121 | BetaCoV/Wuhan/IVDC-HB-05/2019 | China / Hubei Province / Wuhan City | 2019-12-30 | 3 |
| EPI_ISL_402123 | BetaCoV/Wuhan/IPBCAMS-WH-01/2019 | China / Hubei Province / Wuhan City | 2019-12-24 | 4 |
| EPI_ISL_402124 | BetaCoV/Wuhan/WIV04/2019 | China / Hubei Province / Wuhan City | 2019-12-30 | 5 |
| EPI_ISL_402127 | BetaCoV/Wuhan/WIV02/2019 | China / Hubei Province / Wuhan City | 2019-12-30 | 5 |
| EPI_ISL_402128 | BetaCoV/Wuhan/WIV05/2019 | China / Hubei Province / Wuhan City | 2019-12-30 | 5 |
| EPI_ISL_402129 | BetaCoV/Wuhan/WIV06/2019 | China / Hubei Province / Wuhan City | 2019-12-30 | 5 |
| EPI_ISL_402130 | BetaCoV/Wuhan/WIV07/2019 | China / Hubei Province / Wuhan City | 2019-12-30 | 5 |
| EPI_ISL_403928 | BetaCoV/Wuhan/IPBCAMS-WH-05/2020 | China / Hubei Province / Wuhan City | 2020-01-01 | 4 |
| EPI_ISL_403929 | BetaCoV/Wuhan/IPBCAMS-WH-04/2019 | China / Hubei Province / Wuhan City | 2019-12-30 | 4 |
| EPI_ISL_403930 | BetaCoV/Wuhan/IPBCAMS-WH-03/2019 | China / Hubei Province / Wuhan City | 2019-12-30 | 4 |
| EPI_ISL_403931 | BetaCoV/Wuhan/IPBCAMS-WH-02/2019 | China / Hubei Province / Wuhan City | 2019-12-30 | 4 |
| EPI_ISL_402131 | BetaCoV/bat/Yunnan/RaTG13/2013 | China / Yunnan Province / Pu'er City | 2013-07-24 | 6 |
| EPI_ISL_406531 | BetaCoV/Guangdong/20SF013/2037 | China/Guangdong Province | 2020-01-22 | 7 |
| EPI_ISL_406534 | BetaCoV/Guangdong/20SF013/2039 | China/Guangdong Province | 2020-01-22 | 7 |
| EPI_ISL_406535 | BetaCoV/Guangdong/20SF013/2040 | China/Guangdong Province | 2020-01-22 | 7 |
| EPI_ISL_406536 | BetaCoV/Guangdong/20SF013/2041 | China/Guangdong Province | 2020-01-22 | 7 |
| EPI_ISL_406538 | BetaCoV/Guangdong/20SF013/2042 | China/Guangdong Province | 2020-01-23 | 7 |
| EPI_ISL_403932 | BetaCoV/Guangdong/20SF012/2020 | China/Guangdong, China | 2020-01-14 | 7 |
| EPI_ISL_403933 | BetaCoV/Guangdong/20SF013/2020 | China/Guangdong, China | 2020-01-15 | 7 |

| | | | | |
|---|---|---|---|---|
| EPI_ISL_403934 | BetaCoV/Guangdong/20SF013/2021 | China/Guangdong, China | 2020-01-15 | 7 |
| EPI_ISL_403935 | BetaCoV/Guangdong/20SF013/2022 | China/Guangdong, China | 2020-01-15 | 7 |
| EPI_ISL_403936 | BetaCoV/Guangdong/20SF013/2023 | China/Guangdong, China | 2020-01-17 | 7 |
| EPI_ISL_403937 | BetaCoV/Guangdong/20SF013/2024 | China/Guangdong, China | 2020-01-18 | 7 |
| EPI_ISL_406533 | BetaCoV/Guangdong/20SF013/2038 | China/Guangzhou City | 2020-01-22 | 7 |
| EPI_ISL_402132 | BetaCoV/Wuhan/HBCDC-HB-01/2019 | China/Hubei Province | 2019-12-30 | 5 |
| EPI_ISL_406592 | BetaCoV/Guangdong/20SF013/2043 | China/Shenzhen | 2020-01-13 | 8 |
| EPI_ISL_406593 | BetaCoV/Guangdong/20SF013/2044 | China/Shenzhen | 2020-01-13 | 9 |
| EPI_ISL_406594 | BetaCoV/Guangdong/20SF013/2045 | China/Shenzhen | 2020-01-16 | 9 |
| EPI_ISL_406595 | BetaCoV/Guangdong/20SF013/2046 | China/Shenzhen | 2020-01-16 | 9 |
| EPI_ISL_404227 | BetaCoV/Guangdong/20SF013/2027 | China/Zhejiang, China | 2020-01-16 | 10 |
| EPI_ISL_404228 | BetaCoV/Guangdong/20SF013/2028 | China/Zhejiang, China | 2020-01-17 | 10 |
| EPI_ISL_406596 | BetaCoV/France/IDF0372/2020\|EPI_ISL_406596 | France / Ile-de-France / Paris | 2020-01-23 | 11 |
| EPI_ISL_406597 | BetaCoV/France/IDF0373/2020\|EPI_ISL_406597 | France / Ile-de-France / Paris | 2020-01-23 | 11 |
| EPI_ISL_402126 | BetaCoV/Kanagawa/1/2020 | Japan/ Kanagawa Prefecture, Japan | 2020-01-14 | 12 |
| EPI_ISL_406031 | BetaCoV/Guangdong/20SF013/2033 | Taiwan/Kaohsiung City | 2020-01-23 | 13 |
| EPI_ISL_403962 | BetaCoV/Guangdong/20SF013/2025 | Thailand/ Nonthaburi Province | 2020-01-08 | 14 |
| EPI_ISL_403963 | BetaCoV/Guangdong/20SF013/2026 | Thailand/ Nonthaburi Province | 2020-01-13 | 14 |
| EPI_ISL_406223 | BetaCoV/Guangdong/20SF013/2036 | USA / Arizona / Phoenix | 2020-01-22 | 15 |
| EPI_ISL_406034 | BetaCoV/Guangdong/20SF013/2034 | USA / California / Los Angeles | 2020-01-23 | 16 |
| EPI_ISL_406036 | BetaCoV/Guangdong/20SF013/2035 | USA / California / Orange County | 2020-01-22 | 16 |
| EPI_ISL_404253 | BetaCoV/Guangdong/20SF013/2029 | USA / Illinois /Chicago | 2020-01-21 | 17 |
| EPI_ISL_404895 | BetaCoV/Guangdong/20SF013/2030 | USA / Washington / Snohomish County | 2020-01-19 | 18 |

**Citations and acknowledgements to original laboratories generating the data:**

**1 -** National Institute for Communicable Disease Control and Prevention (ICDC) Chinese Center for Disease Control and Prevention (China CDC)
**2 -** The University of Hong Kong - Shenzhen Hospital

**3 -** National Institute for Viral Disease Control and Prevention, China CDC
**4 -** Institute of Pathogen Biology, Chinese Academy of Medical Sciences & Peking Union Medical College
5 - Wuhan Jinyintan Hospital, Wuhan Institute of Virology, Chinese Academy of Sciences
6 - Institute of Pathogen Biology, Chinese Academy of Medical Sciences & Peking Union Medical College
7 - Guangdong Provincial Center for Diseases Control and Prevention; Guangdong Provinical Public Health
8 - Wuhan Jinyintan Hospital, Hubei Provincial Center for Disease Control and Prevention
9 - Shenzhen Key Laboratory of Pathogen and Immunity, National Clinical Research Center for Infectious Disease, Shenzhen Third People's Hospital
10 -Zhejiang Provincial Center for Disease Control and Prevention, Department of Microbiology, Zhejiang Provincial Center for Disease Control and Prevention
11 - Department of Infectious and Tropical Diseases, Bichat Claude Bernard Hospital, Paris, National Reference Center for Viruses of Respiratory Infections, Institut Pasteur, Paris
12 -Deptartment of Virology III, National Institute of Infectious Diseases
13 - Centers for Disease Control, R.O.C. (Taiwan)
14 - Bamrasnaradura Hospital,  Department of Medical Sciences, Ministry of Public Health, Thailand, Thai Red Cross Emerging Infectious Diseases - Health Science Centre, Department of Disease Control, Ministry of Public Health, Thailand
15 - Arizona Department of Health Services, Pathogen Discovery, Respiratory Viruses Branch, Division of Viral Diseases, Centers for Disease Control and Prevention
16 - California Department of Public Health, Pathogen Discovery, Respiratory Viruses Branch, Division of Viral Diseases, Centers for Dieases Control and Prevention
17 - Department of Public Health Chicago Laboratory, Pathogen Discovery, Respiratory Viruses Branch, Division of Viral Diseases, Centers for sControl and Prevention
18 - Providence Regional Medical Center, Division of Viral Diseases, Centers for Disease Control and Prevention

**Supplementary Table 2:** Reference phylogenetic dataset sequences selected for the Genome Detective Coronavirus Tool.

| SARSr-CoV Cluster | Sequence Name | Accession Number | Location | Host |
|---|---|---|---|---|
| Bat SARS-CoV HKU3 | HKU3_3 | DQ084200 | China | Bat |
| Bat SARS-CoV HKU3 | HKU3_6 | GQ153541 | China | Bat |
| Bat SARS-CoV HKU3 | HKU3_9 | GQ153544 | China | Bat |
| Bat SARS-CoV ZXC21/ZC45 | bat_SL_CoVZXC21 | MG772934 | China | Bat |
| Bat SARS-CoV ZXC21/ZC45 | bat-SL-CoVZC45 | MG772933.1 | China | Bat |
| SARS related CoV | As6526 | KY417142 | China | Bat |
| SARS related CoV | BtCoV2732005 | DQ648856 | China | Bat |
| SARS related CoV | BtCoV2792005 | DQ648857 | China | Bat |
| SARS related CoV | Rs3367 | KC881006 | China | Bat |
| SARS related CoV | Rs4237 | KY417147 | China | Bat |
| SARS related CoV | Rs9401 | KY417152 | China | Bat |
| SARS related CoV | WIV1 | KF367457 | China | Bat |
| SARS-CoV Outbreak 2002-3 | BJ182_12 | EU371564 | China | Human |
| SARS-CoV Outbreak 2002-3 | HKU_39849 | AY278491 | Hong Kong | Human |
| SARS-CoV Outbreak 2002-3 | LC4 | AY395001 | China | Human |
| SARS-CoV Outbreak 2002-3 | NC_004718 | NC_004718 | Canada | Human |
| SARS-CoV Outbreak 2002-3 | SARS_ExoN1_2010 | KF514393 | USA | N/A |
| SARS-CoV Outbreak 2002-3 | SARS_WTic_2009 | KF514394 | USA | N/A |
| SARS-CoV Outbreak 2002-3 | SARS_WTic_2010 | KF514388 | USA | N/A |
| SARS-CoV Outbreak 2002-3 | Sin842 | AY559081 | Singapore | Human |
| SARS-CoV Outbreak 2002-3 | Sin848 | AY559085 | Singapore | Human |
| SARS-CoV Outbreak 2002-3 | Sin850 | AY559096 | Singapore | Human |
| SARS-CoV Outbreak 2002-3 | TW7 | AY502930 | Taiwan | Human |
| Wuhan 2019-nCoV | WH-Human1_China_2019-Dec | MN908947 | China | Human |

**Supplementary Table 3:** Evaluation of the Genome Detective Coronavirus Typing Tool to classify coronavirus complete genomes. The classification results were compared to manual phylogenetic analysis . In this table, the following abbreviations are used: TP, total positives; TN, total negatives; FP, false positive; FN, False negative; Sens, sensitivity; Spec, specificity; PPV, positive predicted value; NPVs negative predicted value, ACC, accuracy.

| **Virus species** | **Known** | **TP** | **TN** | **FP** | **FN** | **SENS** | **SPEC** | **ACC** |
|---|---|---|---|---|---|---|---|---|
| Betacoronavirus | 121 | 121 | 311 | 0 | 0 | 100% | 100% | 100% |
| Human Coronavirus HKU1 | 19 | 19 | 413 | 0 | 0 | 100% | 100% | 100% |
| Longquan Rl rat coronavirus | 1 | 0 | 432 | 0 | 1 | 0% | 100% | 100% |
| MERSr-CoV | 97 | 97 | 335 | 0 | 0 | 100% | 100% | 100% |
| Murine Hepatitis Virus | 9 | 9 | 423 | 0 | 0 | 100% | 100% | 100% |
| Rat Coronavirus | 3 | 3 | 429 | 0 | 0 | 100% | 100% | 100% |
| Rousettus bat coronavirus HKU9 | 4 | 4 | 428 | 0 | 0 | 100% | 100% | 100% |
| **SARSr-CoV** | **176** | **176** | **256** | **0** | **0** | **100%** | **100%** | **100%** |
| Tylonycteris bat coronavirus HKU4 | 1 | 1 | 431 | 0 | 0 | 100% | 100% | 100% |
| Zaria_bat_coronavirus | 1 | 0 | 432 | 0 | 1 | 0% | 100% | 100% |
| Total | 432 | | | | | | | |
| **SARSr-CoV Clusters** | **Known** | **TP** | **TN** | **FP** | **FN** | **SENS** | **SPEC** | **ACC** |
| *Bat SARS-CoV HKU3* | 8 | 8 | 424 | 0 | 0 | 100% | 100% | 100% |
| *Bat SARS-CoV ZXC21/ZC45* | 2 | 2 | 430 | 0 | 0 | 100% | 100% | 100% |
| *SARS related CoV* | 6 | 6 | 426 | 0 | 0 | 100% | 100% | 100% |
| *SARSr-CoV outbreak 2002-3* | 112 | 112 | 320 | 0 | 0 | 100% | 100% | 100% |
| ***Wuhan 2019-nCoV*** | **47** | **26** | **406** | **0** | **0** | **100%** | **100%** | **100%** |
| Total | 176 | | | | | | | |
| **Total sequences** | 432 | | | | | | | |

**Supplementary Table 4A:** Nucleotide and protein mutational analysis performed by Genome Detective Coronavirus Typing Tool. This table compares an n2019-CoV query sequence (GenBank Accession Number MN90847) to the reference strain of SARS (NC_004718.3: GenBank). In the table, the following abbreviations are used: Begin, first nucleotide position that reference sequence; End, last nucleotide position that matches the reference sequence; Coverage: coverage of reference sequence genome; Score, nucleotide and amino acid score of AGA; Matches, number of matches; Identities, number of identical nucleotides; I, insertions; D, deletions; M, misaligned; F, Frameshifts; Stop codons, number of stop codons and CDS, coding sequencing.

| **Ref: SARS: NC_004718.3** | **Begin** | **End** | **Coverage** | **Score** | **Concordance** | **Matches** | **Identities** | **I/D** | |
|---|---|---|---|---|---|---|---|---|---|
| **Query: n2019-CoV: MN908947** | 1 | 29677 | 99.8% | 34510 | 58.6% | 29591 (98.9%) | 23621 (79.0%) | 229/86 | |
| CDS | **Begin** | **End** | **Coverage** | **Score** | **Concordance** | **Matches** | **Identities** | **I/D/M/F** | **Stop Codons** |
| 1_orf1ab | 1 | 7074 | 100% | 44034 | 89.0% | 7068 (99.5%) | 6126 (86.2%) | 29/6/0/0 | 1 |
| 2_orf1ab | 1 | 4383 | 100% | 25297 | 83.9% | 4377 (99.2%) | 3553 (80.5%) | 29/6/0/0 | 1 |
| 3_S | 1 | 1256 | 100% | 7176 | 81.6% | 1249 (97.5%) | 976 (76.2%) | 25/7/0/0 | 1 |
| 4_sars3a | 1 | 275 | 100% | 1490 | 75.5% | 275 (99.6%) | 200 (72.5%) | 1/0/0/0 | 1 |
| 5_sars3b | 1 | 155 | 100% | 519 | 52.9% | 155 (99.4%) | 89 (57.1%) | 1/0/0/0 | 5 |
| 6_E | 1 | 77 | 100% | 447 | 96.3% | 76 (98.7%) | 73 (94.8%) | 0/1/0/0 | 1 |
| 7_M | 1 | 222 | 100% | 1419 | 92.9% | 222 (99.6%) | 202 (90.6%) | 1/0/0/0 | 1 |
| 8_sars6 | 1 | 64 | 100% | 300 | 74.6% | 62 (96.9%) | 43 (67.2%) | 0/2/0/0 | 1 |
| 9_sars7a | 1 | 123 | 100% | 758 | 89.3% | 122 (99.2%) | 105 (85.4%) | 0/1/0/0 | 1 |
| 10_sars7b | 1 | 45 | 100% | 252 | 83.7% | 44 (97.8%) | 36 (80.0%) | 0/1/0/0 | 1 |
| 11_sars8a | 1 | 40 | 100% | 100 | 33.8% | 39 (90.7%) | 13 (30.2%) | 3/1/0/0 | 0 |
| 12_sars8b | 1 | 85 | 100% | 26 | 4.2% | 81 (81.8%) | 30 (30.3%) | 14/4/1/1 | 4 |
| 13_N | 1 | 423 | 100% | 2645 | 91.3% | 420 (99.3%) | 383 (90.5%) | 0/3/0/0 | 1 |
| 14_sars9b | 1 | 99 | 100% | 451 | 72.9% | 98 (99.0%) | 72 (72.7%) | 0/1/0/0 | 1 |

**Supplementary Table 4B:** Nucleotide and protein mutational analysis performed by Genome Detective Coronavirus Typing Tool. This table compares an n2019-CoV query sequence (GenBank Accession Number MN90847) to the Bat SARS related CoV sequence, bat_SL_CovZXC21 (MG772934: GenBank). In the table, the following abbreviations are used: Begin, first nucleotide position that reference sequence; End, last nucleotide position that matches the reference sequence; Coverage: coverage of reference sequence genome; Score, nucleotide and amino acid score of AGA; Matches, number of matches; Identities, number of identical nucleotides; I, insertions; D, deletions; M, misaligned; F, Frameshifts; Stop codons, number of stop codons and CDS, coding sequencing.

| **B) bat_SL_CoVZXC21: MG772934** | **Begin** | **End** | **Coverage** | **Score** | **Concordance** | **Matches** | **Identities** | **I/D** | |
|---|---|---|---|---|---|---|---|---|---|
| **n2019-CoV: MN908947** | 1 | 29741 | 99.5% | 44750 | 75.6% | 29638 (99.3%) | 26108 (87.5%) | 182/17 | |
| CDS | **Begin** | **End** | **Coverage** | **Score** | **Concordance** | **Matches** | **Identities** | **I/D/M/F** | **Stop Codons** |
| 1_orf1ab | 1 | 7078 | 99.9% | 47593 | 96.4% | 7071 (99.6%) | 6767 (95.3%) | 28/1/6/0 | 1 |
| 2_orf1ab | 1 | 4387 | 99.9% | 28933 | 96.3% | 4380 (99.3%) | 4197 (95.2%) | 28/1/6/0 | 1 |
| 3_S | 1 | 1253 | 99.4% | 7104 | 83.8% | 1243 (97.6%) | 1015 (79.7%) | 29/2/10/0 | 1 |
| 4_sars3a | 1 | 276 | 100% | 1847 | 93.7% | 276 (100%) | 254 (92.0%) | 0/0/0/0 | 1 |
| 5_sars3b | 1 | 156 | 100% | 755 | 75.1% | 156 (100%) | 124 (79.5%) | 0/0/0/0 | 5 |
| 6_E | 1 | 77 | 98.7% | 474 | 100% | 76 (100%) | 76 (100%) | 0/0/0/0 | 1 |
| 7_M | 1 | 223 | 100% | 1521 | 98.6% | 223 (100%) | 220 (98.7%) | 0/0/0/0 | 1 |
| 8_sars6 | 1 | 64 | 96.9% | 369 | 86.6% | 62 (100%) | 58 (93.5%) | 0/0/0/0 | 1 |
| 9_sars7a | 1 | 123 | 99.2% | 776 | 91.0% | 122 (100%) | 108 (88.5%) | 0/0/0/0 | 1 |
| 10_sars7b | 1 | 45 | 97.8% | 288 | 93.8% | 44 (100%) | 41 (93.2%) | 0/0/0/0 | 1 |
| 11_sars8a | 1 | 43 | 97.7% | 331 | 98.5% | 42 (100%) | 40 (95.2%) | 0/0/0/0 | 0 |
| 12_sars8b | 1 | 99 | 96.0% | 453 | 74.0% | 95 (100%) | 73 (76.8%) | 0/0/2/0 | 4 |
| 13_N | 1 | 422 | 99.5% | 2707 | 94.6% | 419 (99.5%) | 396 (94.1%) | 1/1/0/0 | 1 |
| 14_sars9b | 1 | 99 | 99.0% | 446 | 72.3% | 98 (100%) | 72 (73.5%) | 0/0/0/0 | 1 |

**Supplementary Table 4C:** Nucleotide and protein mutational analysis performed by Genome Detective Coronavirus Typing Tool. This table compares an n2019-CoV query sequence, BetaCoV/France/IDF0373/2020|EPI_ISL_406597, isolated in France, Jan 2010, (GISAID Accession Number ISL_406597) to the first n2019-CoV sequence, isolated in Wuhan, Dec 2019 (MN908947: GenBank). The following abbreviations are used: Begin, first nucleotide position that reference sequence; End, last nucleotide position that matches the reference sequence; Coverage: coverage of reference sequence genome; Score, nucleotide and amino acid score of AGA; Matches, number of matches; Identities, number of identical nucleotides; I, insertions; D, deletions; M, misaligned; F, Frameshifts; Stop codons, number of stop codons and CDS, coding sequencing.

| **C) n2019-CoV: MN908947** | **Begin** | **End** | **Coverage** | **Score** | **Concordance** | **Matches** | **Identities** | **I/D** | |
|---|---|---|---|---|---|---|---|---|---|
| n2019-CoV: BetaCoVFrance IDF03732020 | 20 | 29914 | 99.4% | 59606 | 99.9% | **29809 (100%)** | **29806 (99.9%)** | 0/0 | |
| CDS | **Begin** | **End** | **Coverage** | **Score** | **Concordance** | **Matches** | **Identities** | **I/D/M/F** | **Stop Codons** |
| 1_orf1ab | 1 | 7103 | 99.9% | 49596 | 100% | 7097 (100%) | 7097 (100%) | 0/0/0/0 | 1 |
| 2_orf1ab | 1 | 4412 | 99.9% | 30277 | 100% | 4406 (100%) | 4406 (100%) | 0/0/0/0 | 1 |
| 3_S | 1 | 1281 | 99.5% | 8985 | 99.9% | **1274 (100%)** | **1273 (99.9%)** | 0/0/0/0 | 1 |
| 4_sars3a | 1 | 276 | 100% | 1944 | 99.4% | **276 (100%)** | **275 (99.6%)** | 0/0/0/0 | 1 |
| 5_sars3b | 1 | 156 | 100% | 964 | 99.6% | **156 (100%)** | **155 (99.4%)** | 0/0/0/0 | 5 |
| 6_E | 1 | 77 | 98.7% | 474 | 100% | 76 (100%) | 76 (100%) | 0/0/0/0 | 1 |
| 7_M | 1 | 223 | 100% | 1538 | 100% | 223 (100%) | 223 (100%) | 0/0/0/0 | 1 |
| 8_sars6 | 1 | 64 | 96.9% | 400 | 100% | 62 (100%) | 62 (100%) | 0/0/0/0 | 1 |
| 9_sars7a | 1 | 123 | 99.2% | 846 | 100% | 122 (100%) | 122 (100%) | 0/0/0/0 | 1 |
| 10_sars7b | 1 | 45 | 97.8% | 316 | 100% | 44 (100%) | 44 (100%) | 0/0/0/0 | 1 |
| 11_sars8a | 1 | 43 | 97.7% | 339 | 100% | 42 (100%) | 42 (100%) | 0/0/0/0 | 0 |
| 12_sars8b | 1 | 99 | 96.0% | 608 | 100% | 95 (100%) | 95 (100%) | 0/0/2/0 | 4 |
| 13_N | 1 | 423 | 99.3% | 2860 | 99.9% | **420 (100%)** | **419 (99.8%)** | 0/0/0/0 | 1 |
| 14_sars9b | 1 | 99 | 99.0% | 618 | 100% | 98 (100%) | 98 (100%) | 0/0/0/0 | 1 |

**Supplementary Table 4C-II:** This table compares an n2019-CoV query sequence, BetaCoV/France/IDF0373/2020|EPI_ISL_406597, isolated in France, Jan 2010, (GISAID Accession Number ISL_406597) to the first n2019-CoV sequence, isolated in Wuhan, Dec 2019 (MN908947: GenBank). The French isolated sequence has two nucleotide (NT) mutations at genome position 22551 (G to T) and 26016 (G to T). These mutations affect three proteins: E2 glycoprotein (Uniprot Accession Number: NP_828851.1), Hypothetical protein sars3a (Uniprot Accession Number: NP_828852.2) and Hypothetical protein sars3b ((Uniprot Accession Number: NP_828853.1). Please note that NT mutation 26016G>T affects two open reading frames that code two hypothetical proteins, sars3a and sars3b.

| Mutations NT | **22551G>T, 26016G>T** |
|---|---|
| **Proteins** | **Mutations:** |
| E2 glyco (NP_828851.1 ) | Protein mutations:<br>V354F (22551G>T)<br>Codon mutations:<br>GTC354TTC (22551G>T) |
| sars3a (NP_828852.2) | Protein mutations:<br>G250V (26016G>T)<br>Codon mutations:<br>GGT250GTT (26016G>T) |
| sars3b (NP_828853.1) | Protein mutations:<br>V110F (26016G>T)<br>Codon mutations:<br>GTT110TTT (26016G>T) |
